# Fano factor as a key measure of sensitivity in biological networks

**DOI:** 10.1101/2025.04.03.646967

**Authors:** Kinshuk Banerjee, Biswajit Das, Pintu Patra

**Affiliations:** Department of Chemistry, Acharya Jagadish Chandra Bose College, 1/1B A. J. C. Bose Road, Kolkata 700 020, India; School of Artificial Intelligence (AI), Amrita Vishwa Vidyapeetham (Amrita University), Amritanagar, Ettimadai, Coimbatore, Tamil Nadu 641112, India; Department of Physics, Indian Institute of Technology, Kharagpur, India

**Keywords:** Cooperative binding networks, Hill slope, Fano factor

## Abstract

Sensitivity of a biological process is vital for its proper function and regulation. Biological networks in equilibrium display such sensitivity mainly through cooperative binding of ligands to network elements comprised of multiple sites of an oligomeric protein. The classical measure of binding cooperativity is the Hill slope obtained empirically from binding isotherm data, known as the Hill plot. Hill slope value greater (less) than unity indicates positive (negative) cooperativity. In this study, we have shown that the Hill slope is actually a combination of two Fano factors associated with the number of bound and empty sites of the network, respectively. Taking a ladder network capable of showing positive cooperativity only, we have analytically proved that if the variation of Fano factors with fractional saturation of the network is non-linear, the binding is cooperative and if the variation is linear, the binding is non-cooperative with Hill slope equal to unity. Further, through numerical analyses of a linear network that can exhibit both positive and negative cooperativity, it is established that the curves of Fano factor for the positive (negative) cooperative cases reside above (below) the line representing the non-cooperative case. Our results, thus, confirm the role of Fano factor as the crucial measure of sensitivity in biological networks.

## 1 Introduction

In cellular biology, the sensitivity of a biophysical process is a key factor characterized by the experimentally determined response of that process[1, 2]. Generally, a biophysical process is associated with an optimal and adaptive network that has evolved to exhibit the diverse response[3, 4]. A cellular network is formed with various biological components, called network elements, which can be of similar or different types of cells or cellular entities including proteins, substrates, amino acids, etc. Importantly, elements of a cellular network are dynamic in nature as they are modified by binding and unbinding of the signaling molecules or ligands to them. Thus, the response of a network not only characterizes the sensitivity of the biophysical process but also inherently carries statistical and dynamical information about the network elements[5]. Different cooperative binding processes of the ligands to a network formed by the multiple binding sites of a macromolecule is a fundamental mechanism for sensitivity in biological systems[6–8]. The traditional measure of such sensitivity is the phenomenological Hill slope[9], which is determined from the Hill plot generated using the experimentally measurable response quantity, *viz*., the fractional saturation, *θ*(0 ≤ *θ* ≤ 1) of the network as a function of the ligand concentration, [*L*]. Mathematically, the Hill slope is defined as the logarithmic sensitivity of the function 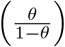 with respect to [*L*] *i*.*e*., 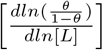 [10]. The value of the Hill slope at the half-saturation point (*θ* = 0.5), denoted as the Hill coefficient[11], characterizes the nature of cooperative binding of the ligands. A Hill coefficient value greater than unity indicates that the ligands bind to the network elements in a positive cooperative manner. Consequently, the network exhibits the delayed sigmoidal response[9]. A value of the Hill coefficient equal to unity indicates noncooperative binding of the ligand where the network shows a rectangular hyperbolic response[10, 11]. When the Hill coefficient becomes less than unity, the binding of ligands occurs with negative cooperativity.

The binding of ligands to network elements is basically a stochastic process at cellular level[12, 13]. The number of bound (empty) elements, say *n* (*n*_*T*_ −*n*), thus becomes a random variable with a probability distribution where *n*_*T*_ is the total number of network elements. Theoretically, the distribution can be estimated by modeling the ligand binding dynamics as a random walk problem solved with the chemical master equation technique[12]. Interestingly, such a distribution can be expressed in a general form for the various networks used to theoretically model cooperative binding [10, 11]. The Hill slope is then shown to be given as the ratio of the variances of the binding number in the cooperative case to that of the corresponding non-cooperative case[11]. Now, the strategic combination of both *θ* and (1 − *θ*) in the Hill slope, although quite successful in the characterization of cooperativity, involves apparently redundant usage of the same quantity, *θ*. This fact, along with the empirical nature of the Hill slope, motivates us to look for a theoretical measure of sensitivity with judicial choice of network variables. As the Hill slope is a ratio of variances, it will be interesting to explore the issue of sensitivity in terms of the dispersion of *n* and (*n*_*T*_ −*n*) which, interestingly, have the same variance *σ*^2^ = ⟨*n*^2^ ⟩− ⟨*n* ^2^. ⟩

In view of the discussion above, we investigate the Fano factor, defined as the ratio of variance to mean, with respect to both the number of bound and empty sites. The importance of Fano factor in the quantification of noise in various biological networks are well-established, *e*.*g*., in gene expression and regulatory networks[14–19]. The Fano factor has also been shown to follow a universal bound, revealing important information on enzyme kinetic networks[20]. In this context, we ask three questions: (i) *What is the relation, if any, between the Hill slope and the Fano factors associated with the two random variables n and* (*n*_*T*_ −*n*)*?* (ii) *Is it possible to replace the role of Hill coefficient in terms of these Fano factors?* (iii) *Does one really need two such Fano factors to characterize the nature of sensitivity?* The last question arises from the fact that *n* and (*n*_*T*_ − *n*) are, obviously, not independent for a given network size *n*_*T*_, as are *θ* and (1 − *θ*) in the classical Hill plot.

In this study, we have shown that the Hill slope can be expressed as a combination of Fano factors that indicates the crucial role of the latter in describing the response of biological networks. Next, it is further established that the Fano factors can individually provide the same information regarding the nature of sensitivity as given by the Hill coefficient. Thus, these features make the Fano factor a fundamental statistical measure of sensitivity.

## 2 Hill slope as a combination of Fano factors

The Hill slope is determined in the form of logarithmic sensitivity from the Hill plot as [9, 11, 13, 21–23]

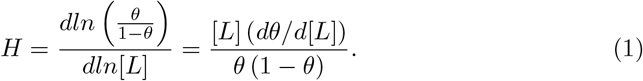

Here 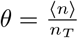, where ⟨*n*⟩ is the mean binding number of the network elements. Statistically, the Hill-slope is shown to be related to the variance of the binding number *σ*^2^ as[9, 13]

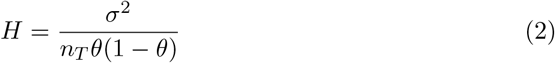

where the denominator in the right-hand-side is the variance of the corresponding non-cooperative case. The variance of the cooperative case, *σ*^2^, becomes higher or lower than that of the non-cooperative one for positive and negative cooperativity, respectively, with the mean binding number being the same in both scenarios for a given ligand concentration[11].

Interestingly, the statistical expression of Hill slope given in Eq.(2) can be split into two Fano factors as the ratio of variance to mean:

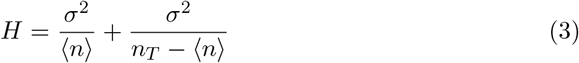

The first term in the r.h.s. of Eq.(3) is the Fano factor in terms of the bound-state random variable *n*; the second term then represents the Fano factor related to the unbound or vacant state random number (*n*_*T*_ − *n*) with mean (*n*_*T*_ − ⟨*n*⟩) and the same *σ*^2^. Eq.(3) shows that the Hill slope is actually a derived quantity from these two Fano factors.

## 3 Fano factor as an indicator of cooperative response

The nature of the variation of *θ* with ligand concentration [*L*] provides qualitative information on whether the underlying binding mechanism is cooperative or not. The Hill plot and, subsequently, the Hill coefficient derived from it, then quantifies the cooperativity. Now, it has already been shown in Sec.2, that the Hill slope comes from Fano factors. Hence, it would be pertinent to investigate whether the Fano factor can play the same role as the Hill coefficient. To this end, we explore the relation of Fano factor with fractional saturation *θ* for two seminal models of cooperativity, *viz*. the Monod-Wyman-Changeux (MWC) network[6] and the Koshland-Nemethy-Filmer (KNF) network[7]. Here, we denote the MWC network as the ladder network and the KNF network as the linear network. Details of these networks are given in Fig.1.

**Fig. 1.**
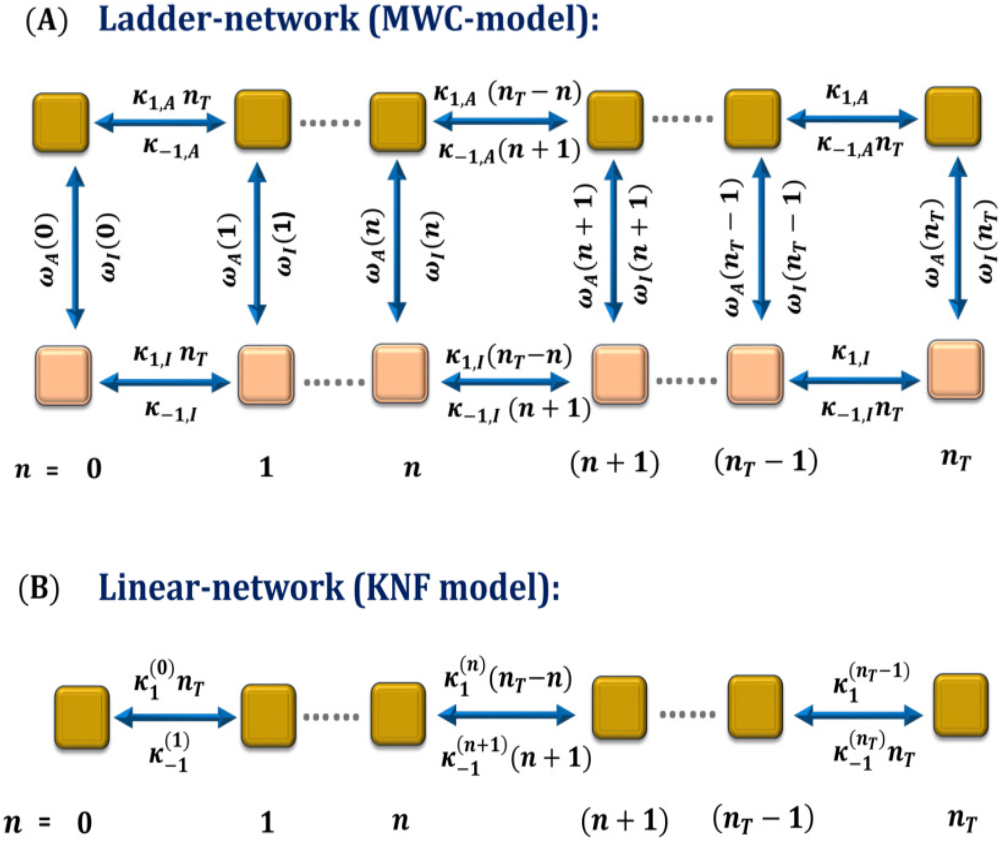
Schematic diagrams of (A) a ladder network akin to the Monod-Wyman-Changeux (MWC) model and (B) a linear network equivalent to the Koshland-Nemethy-Filmer (KNF) model. In (A), each network element remains either in inactive conformational state, ‘I’, portrayed here as light orange square and active conformational state, ‘A’, depicted as deep gold square. Here, ‘*n*’ denotes the number of network elements being bound with ligand molecules. The A-conformation has a ligand-binding affinity greater than that of the I-conformation. In the absence of ligands, the network element resides in I-state. Due to gradual binding of the ligands to the elements of the I-state, the population of the corresponding state increases and suddenly, I-state switches to the A-conformational state giving the sigmoidal response. (B) In the linear (KNF) network, it is considered that when a ligand binds to the inactive conformation of the network elements, concomitantly, the network elements switch to the active conformation. In this model, a network element can not reside in active or inactive state. Moreover, in the KNF network it is considered that the ligand binding affinity from one network element to other may differ. Furthermore, in both of these models, it is considered that binding of one ligand to a network element is sufficient to make transition from one element to another. The details of the transition probabilities in forward and backward directions are also given (see text).

### 3.1 Fano factor in terms of fractional saturation: MWC network

The ladder-like MWC network [6] is shown schematically in Fig.1(A). The network kinetics is described in terms of the number of ligand-bound elements in the state-*χ*(= *A, I*) with 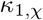 and 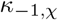 being the forward and backward rate constants. The rate constant for the conformational transitions from *A* to *I* is denoted by *ω*_*A*_(*n*) and that of the opposite transition by *ω*_*I*_ (*n*) for the *n*-th binding state. Further, the equilibrium constants are defined as

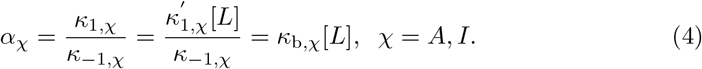

Here *α*_*I*_ = *cα*_*A*_ and 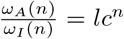 with 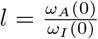. For significant cooperative behavior, *l* ≫1, *c* ≪1 [10]. Now, the number of bound elements is a random variable and the corresponding probability distribution can be determined by solving the chemical master equation (CME) for the network given by

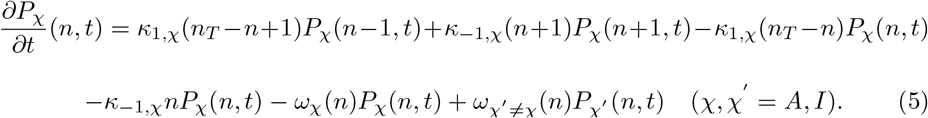

Here, *P*_*χ*_(*n, t*) denotes the probability that the multimeric protein resides in conformation *χ* with *n*-number of binding sites. The equilibrium solution for this network becomes

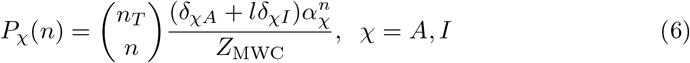

where 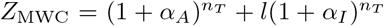 . Summing over the conformation index *χ*, one can express the equilibrium distribution as

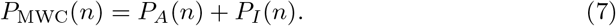

Using the above distribution, the corresponding mean and variance can be determined (see appendixA for details of the calculation).

Now, the variance *σ*^2^ can be generally written as

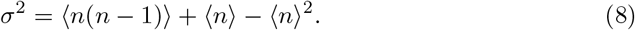

Then, the Fano factor *σ*^2^*/*⟨*n*⟩ becomes

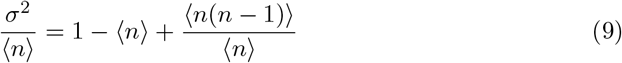

The expressions of ⟨*n*⟩ and ⟨*n* (*n* − 1) ⟩ for the MWC network are as follows (for details, see appendixA):

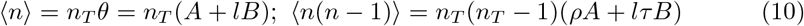

where

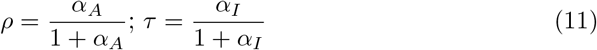

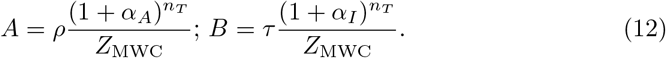

Now, for *α*_*A*_ = *α*_*I*_ with *c* = 1, from Eq.(11) we have *ρ* = *τ* . Then, it follows from Eq.(10) and Eq.(12) that

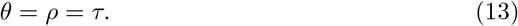

The above form of *θ* actually corresponds to a case with no cooperativity. See Appendix B for details. We would like to mention here that the MWC network does not allow negative cooperativity.

Using equations (10-12), one can express *A* and *B* as a function of *θ, ρ, τ* as

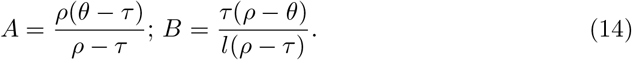

Then, from Eq(9), Eq(10) and Eq.(14), we finally get (see appendixC for details)

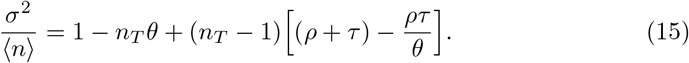

Eq.(15) clearly indicates that *the Fano factor can vary non-linearly with θ* except for the following two cases: (i) when *n*_*T*_ = 1 and (ii) when *ρ* = *τ* = *θ*. In both the scenarios, the Fano factor varies linearly with *θ* given by

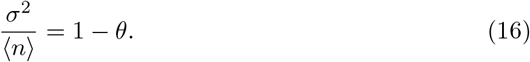

Now, both of the above cases have no scope of ultrasensitivity or positive cooperativity. This yields the main result of this study: *the Fano factor* 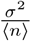 *varies non-linearly with θ for the cooperative case (c* ≠ 1*) whereas the variation becomes linear when the binding is non-cooperative (c* = 1 *or n*_*T*_ = 1). Fig.2 confirms this statement for different values of parameter *c*. It is also clear that the other Fano factor, 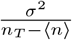, is not required for this type of characterization. However, it also independently carries the same information.

**Fig. 2.**
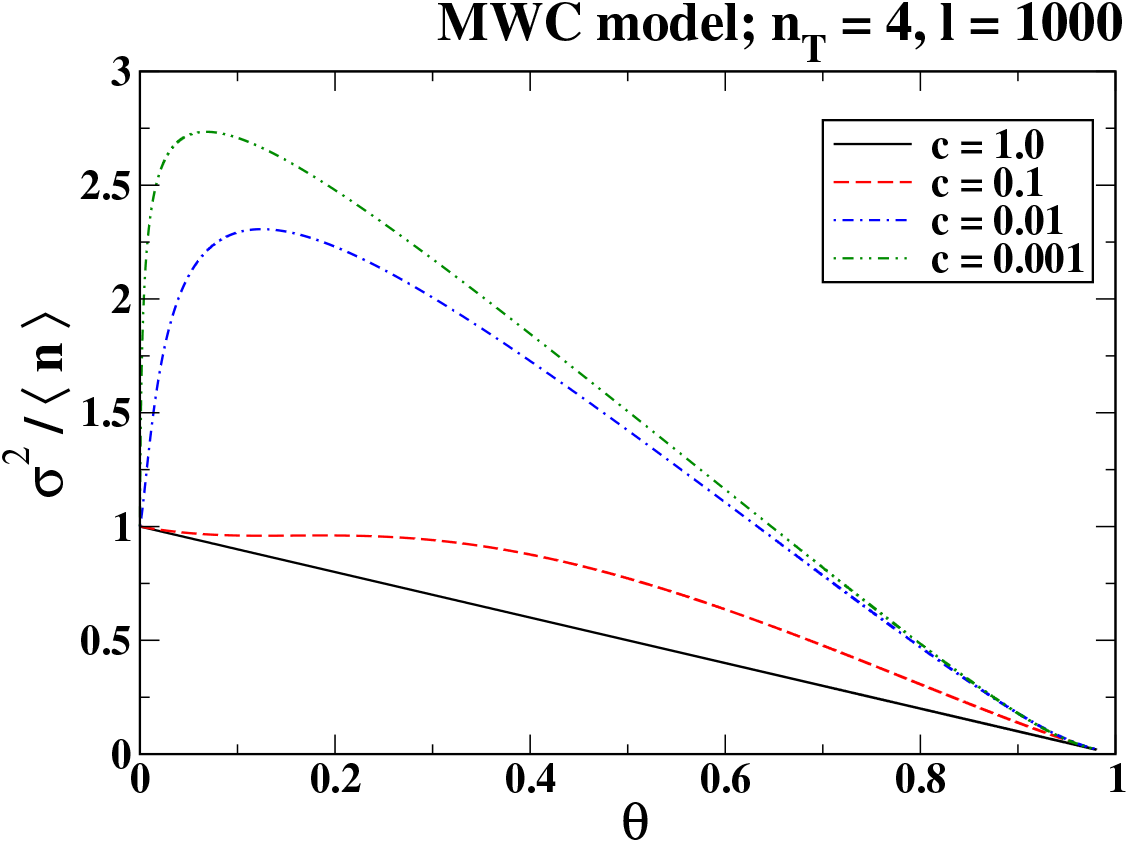
Variation of the Fano factor with fractional saturation *θ* for the MWC model with *n*_*T*_ = 4, *l* = 1000. The case with *c* = 1.0 gives the non-cooperative binding where the Fano factor varies linearly. Cases with *c <* 1.0 show increasing positive deviation with decreasing *c* indicating higher positive cooperativity.

### 3.2 Limiting Cases

We now consider some limiting values of the Fano factor at low ligand concentration [*L*] with *α*_*A*_, *α*_*I*_ *<<* ≤ 1. The parameter values are taken as follows: *l >>* 1, *c* 1, ensuring significant positive cooperativity.

**Case 1:** *lc >>* 1, *lc*^2^ *>>* 1

It follows from Eqs.(10-12) and Eq.(15) that the Fano factor at low [*L*] can be expressed as

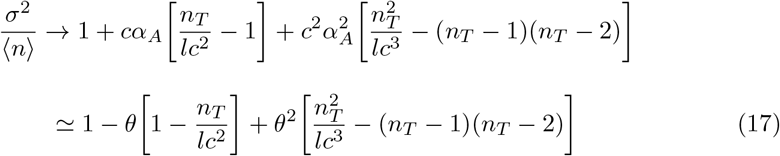

as it follows from Eq.(10-12) that *θ* ≈ *cα*_*A*_ at low [*L*]. As *lc*^2^ *>>* 1, it follows from Eq.(17) that the Fano factor can be less than 1 (one) at low [*L*] (and hence low *θ*). This is also evident in Fig.2 for the case with *l* = 1000, *c* = 0.1, *lc*^2^ = 10 (red dashed curve). In this case, the Fano factor thus approaches the limiting value of unity from below as a function of *θ*.

**Case 2:** *lc >>* 1, *lc*^2^ *<<* 1

In this case, again with *θ* ≈ *cα*_*A*_ at low [*L*], one has

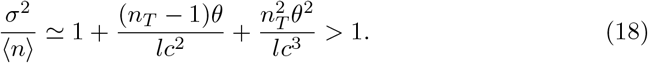

Thus, here the Fano factor dose not become less than unity and rises sharply with *θ* as *lc*^2^ *<<* 1. Again, this is clear from Fig.2 for the case with *l* = 1000, *c* = 0.01, *lc*^2^ = 0.1 (blue dash-dot curve). In this case, the Fano factor approaches the limiting value of unity from above as a function of *θ*.

For the sake of completeness, one can easily find from Eq.(15) that at saturating [*L*] with 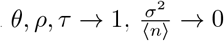. This fact is again clear from Fig.2.

### 3.3 The KNF network

The KNF network[7] is shown in Fig.1(B) schematically. It can be visualized as the linear part of a ladder network with only one kind of sites depicted in Fig.1(A). In this network, the ligand molecules bind or dissociate from the element of a network sequentially with state-dependent equilibrium constants. This network is capable of showing both positive and negative cooperativity. The details of the CMEs for this network to find the probability distribution are given in AppendixD. The equilibrium solution is given as

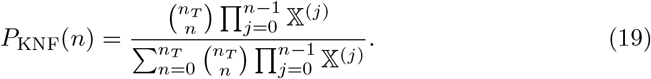

In Eq.(19), the state-dependent binding constants are defined as

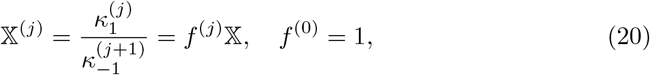

where

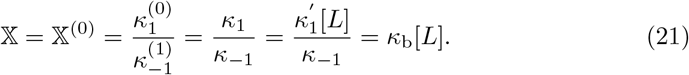

For *f* ^(*j*)^ = *f* = 1 ∀*j*, the KNF network reduces to a non-cooperative (NC) network. With *f >* 1, the network shows positive cooperativity whereas for *f <* 1, negative cooperative behavior is displayed.

The variation of the Fano factor with *θ* for different parameter values is depicted in Fig.3. For *f* = 1.0, the NC scenario gives the straight line. Different cooperative cases (*f* ≠ 1.0) produce non-linear curves, as expected. In case of positive cooperativity, the curves lie above the line of the NC case, whereas for negative cooperativity, the curves reside below that line. These features are quite evident from Fig.3. Thus, the Fano factor can successfully discriminate between positive and negative cooperative cases.

**Fig. 3.**
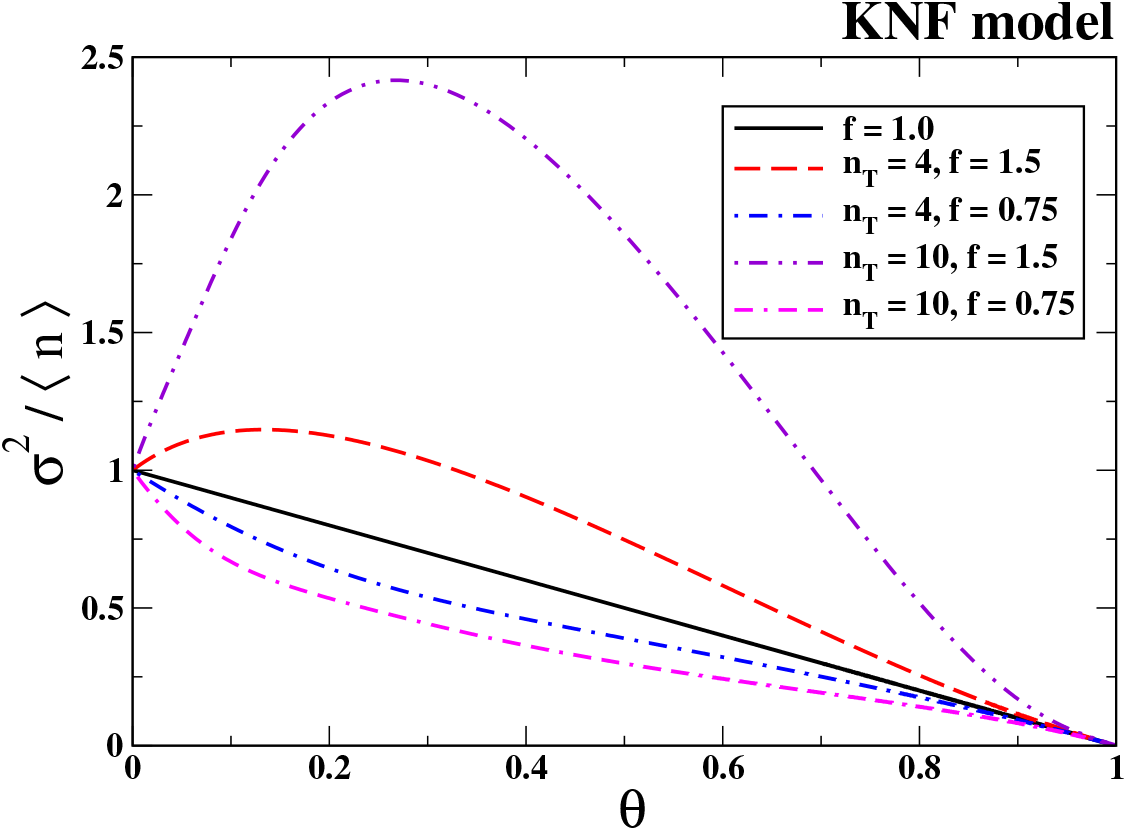
Variation of the Fano factor with fractional saturation *θ* for the KNF model. The case with *f* = 1.0 gives the non-cooperative scenario where the Fano factor follows a linear trend. Cases with *f >* 1.0, specifically *f* = 1.5 here, shows positive deviation from linearity indicating positive cooperativity. On the other hand, cases with *f <* 1.0, specifically *f* = 0.75 here, shows negative deviation indicating negative cooperativity. This correspondence between deviation of the Fano factor from linearity with cooperativity is shown for two system sizes, *n*_*T*_ = 4 and *n*_*T*_ = 10.

## 4 Discussion and Conclusion

The Hill slope and the corresponding Hill coefficient act as the classical measure of sensitivity in biological networks with cooperative ligand binding to the network elements. The Hill plot is constructed from a function of the fractional saturation, *θ* and the fractional unsaturation, (1 − *θ*), of the network. This type of combination characterizes the nature of cooperative response and also quantifies it in terms of the Hill coefficient. However, employing both *θ* and (1 − *θ*) in this phenomenological measure, although quite successful, begs the question whether such usage is redundant and more fundamental measures of sensitivity exist. The theoretical basis for the search of such a measure comes from the fact that the Hill slope can be represented as the ratio of the variance of the number of bound network elements, *n*, in the cooperative case to that in the corresponding non-cooperative case[11]. Hence, we looked for a statistical measure related to the dispersion of the observable response of the network. Now, the Fano factor, defined as the ratio of variance to mean, is such a candidate, and it is an established measure of noise in various networks [14–19]. So, we explored this measure as an educated guess.

We have shown that the Fano factor associated with the number of bound network elements, denoted by *σ*^2^*/* ⟨*n*⟩, and its counterpart in terms of the number of empty elements, *σ*^2^*/*(*n*_*T*_ − ⟨*n*⟩), combine to form the Hill slope as given in Eq.(3), *n*_*T*_ being the total number of binding sites. This result puts the Fano factor on a strong theoretical footing as a fundamental measure of sensitivity. As the random variables *n* and (*n*_*T*_ − *n*) are not independent for a given network size, then comes the acid test: can, out of these two Fano factors, any one independently characterize the nature of sensitivity like the Hill coefficient? Exploring the ladder network first, capable of showing positive cooperativity only, we have analytically established that the nature of variation of the Fano factor *σ*^2^*/* ⟨*n*⟩ as a function of *θ* serves that purpose as follows: *if the variation of σ*^2^*/* ⟨*n*⟩ *with θ is nonlinear, the network is cooperative, and if the variation is linear, then the network is noncooperative*. Further, for the ladder network, the Fano factor curves for the cooperative cases never come below the line representing the noncooperative case. This feature indicates a positive cooperativity.

Now, from the discussion in the previous paragraph, one expects that for a network that allows both positive and negative cooperative behaviors, the curves of the Fano factor would lie both above and below the non-cooperativity line, respectively. We have shown this indeed to be true by numerically exploring a linear network with state-dependent equilibrium constants. This network can exhibit both positive and negative cooperativity depending on whether the binding affinity successively rises or falls along the network, respectively. For the positive cooperative case, the curves lie above the line of the non-cooperative case; in case of negative cooperativity, the curves stay below that line. Hence, non-linear variation of the Fano factor as a function of fractional saturation indicates cooperativity, and the nature of the latter can be revealed through the relative position of the Fano factor curve with respect to the line of the non-cooperative case. We further point out that this line corresponding to non-cooperative binding can be very easily constructed as it starts from unity and falls to zero with a unit slope. This would allow one to easily interpret the relevant experimental data to determine the presence and nature of cooperative response.

Our results in this study clearly establish the Fano factor as a vital statistical measure of sensitivity in biological networks in equilibrium, where cooperative binding of ligands plays a major role in the development of such sensitivity. The criterion of cooperativity has been concisely given in terms of any one of the two Fano factors, and the quantification is also satisfactory. As the Hill slope is shown to be basically a combination of the two Fano factors, the latter serves as a more fundamental measure of sensitivity. The Fano factor yields the same conclusions about nature and degree of cooperativity as obtained from the Hill plot and the Hill coefficient. Therefore, this study confirms the Fano factor as a crucial statistical measure of sensitivity that gives a deeper theoretical basis for the classical phenomenological measures. The applicability of this measure will be tested in more general networks under nonequilibrium conditions in the future.

## Appendix A Mean and variance of the MWC network

The probability distribution derived in Eq.(7) of the main text is employed to determine the moment generating function to compute all the moments defined below

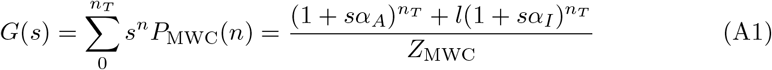

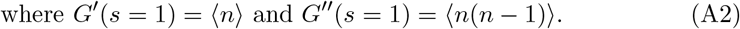

The first moment or mean is then given as

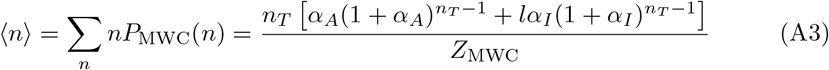

where 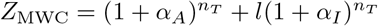 . Using similar approach one can compute

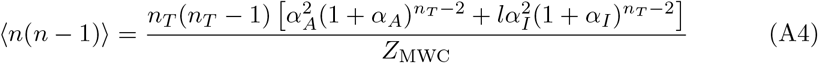

Now, we denote

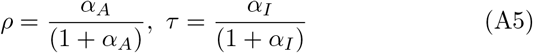

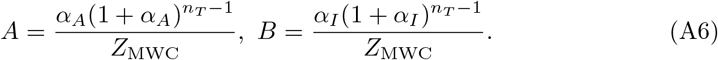

Therefore

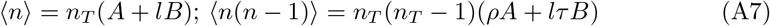

and

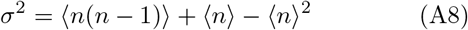

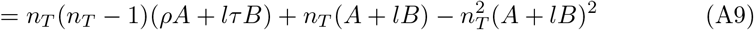

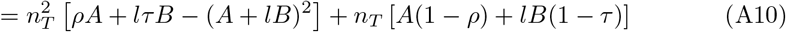

Now we will solve for the term in square brackets, the second term is

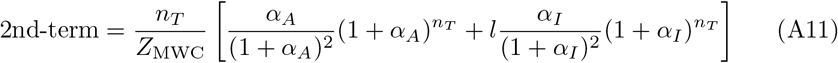

The first term requires major simplification (discussed later) and leads to

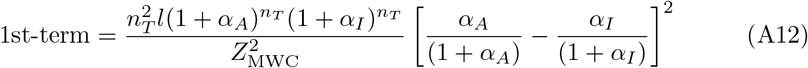

Together, it leads to

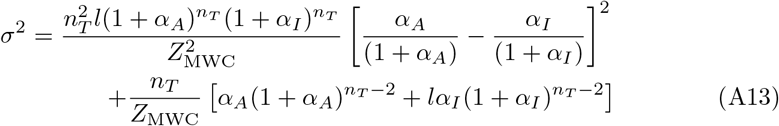

### A.1 First term simplification

The first term is written as

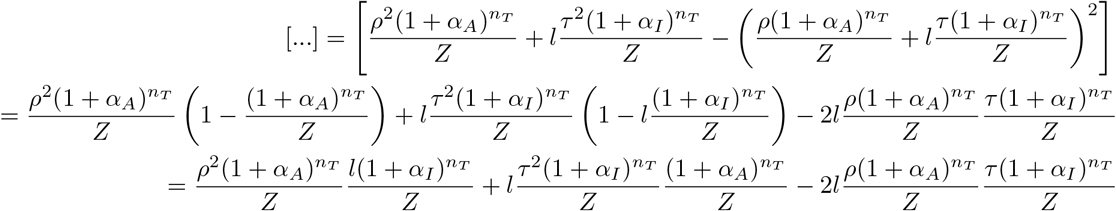

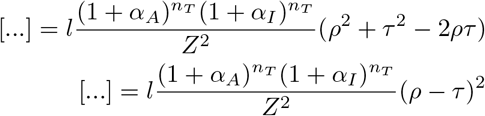

## Appendix B Detailed expressions of the two Fano factors: MWC network

Now, from Eq.(A3) and Eq.(A13), we get the exact expression of Fano factor as

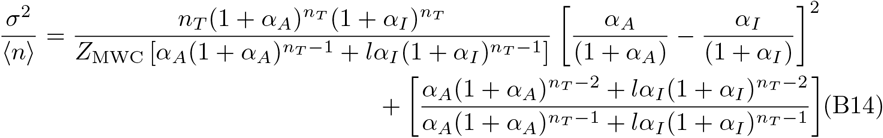

For *α*_*R*_ = *α*_*T*_ = *α, i*.*e*., *c* = 1, the system becomes non-cooperative (NC). The mean is then given as

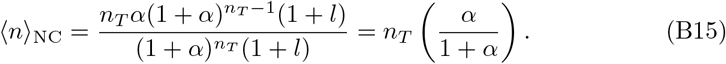

Eq.(B15) is the same as that of Eq.(13) of the main text, as should be the case. Now, for *c* = 1, the first term in the right-hand-side of Eq.(B14) vanishes and the Fano factor reduces to

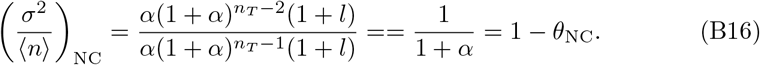

Similarly, one can write the other Fano factor in Eq.(3) as

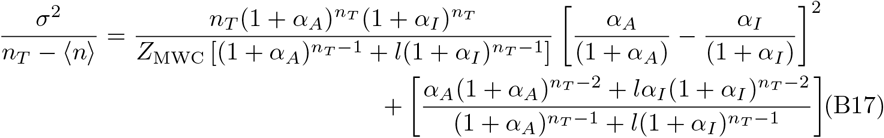

where

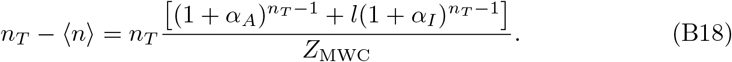

For the NC case, this Fano factor becomes

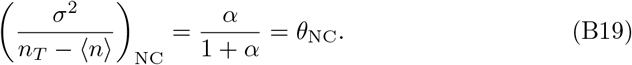

Hence, following Eq.(3) of the main text, the Hill slope becomes

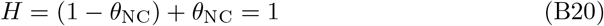

as should be the case for the NC scheme.

## Appendix C Fano factor in terms of fractional saturation: MWC network

The Fano factor can be expressed as

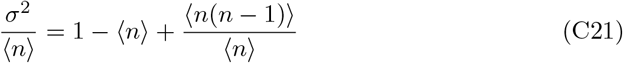

As already mentioned in AppendixA, we denote

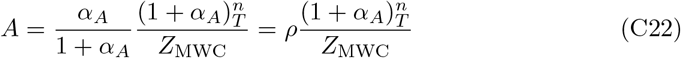

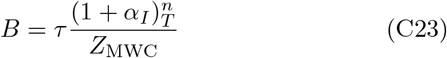

Then we have the following relations

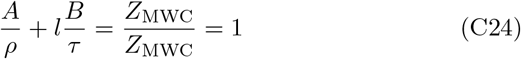

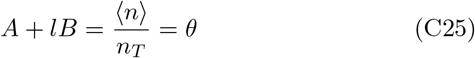

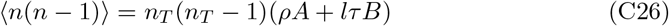

Therefore

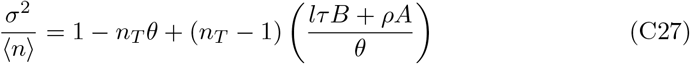

We need A and B in terms of *θ*. We can write *A* = *θ* − *lB*; then

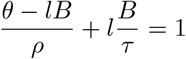

that gives

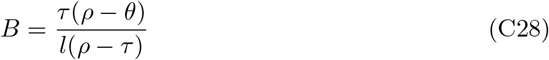

Then

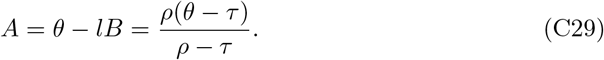

Now, after some algebra, we have

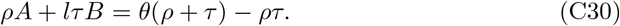

Finally, this produces Eq.(15) of the main text as

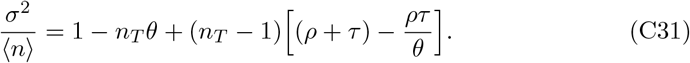

## Appendix D The KNF network

The chemical master equation for this network can be generally formulated as

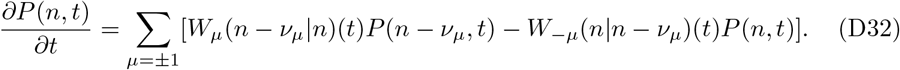

Here, *µ* is the reaction index with *µ* = 1 for the forward reaction and *µ* = −1 for the reverse reaction. The stoichiometric coefficients *ν*_*µ*_ have values *ν*_1_ = 1 and *ν*_−1_ = − 1. The probability of having ‘*n*’ number of elements that are occupied by ligand molecules at time *t* is denoted by *P* (*n, t*) with *n* running from 0 to *n*_*T*_, the total number of elements in the network. The transition probabilities between the elements, *W*_*µ*_, are defined for the KNF network by

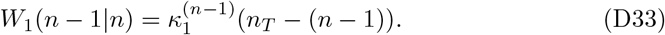

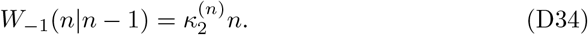

For clarity, we label *P* (*n, t*) by *P*_KNF_(*n, t*) for the KNF network. The equilibrium solution of the CMEs for the above networks is obtained by setting the right-hand side of the set of equations, Eq.(D32), equal to zero. It is given below:

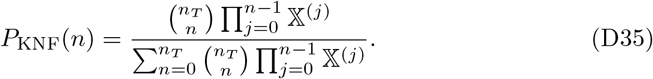

The state-dependent binding constants are defined as

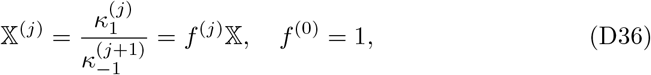

where

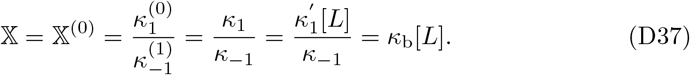

For 𝕏^(*j*)^ = 𝕏 ∀*j*, the KNF network reduces to a non-cooperative (NC) network with a binomial distribution

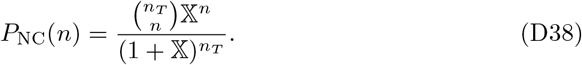

The mean binding number is defined as ⟨*n*⟩ = Σ_n_*nP* (*n*). However, it can be analytically determined in a closed form only for the NC scheme given by

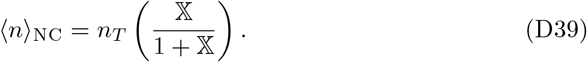

